# Stromal SRD5A2 promotes prostate growth through WNT5A-LEF1-IGF1 signaling in benign prostatic hyperplasia

**DOI:** 10.1101/2025.05.22.655540

**Authors:** Christina Sharkey, Boqing Gu, Xingbo Long, Yao Tang, Nicolas Patsatzis, Steven Li, Aria F. Olumi, Zongwei Wang

## Abstract

Steroid 5α-reductase type 2 (SRD5A2) is a key enzyme in androgen metabolism and a pharmacologic target in benign prostatic hyperplasia (BPH). While SRD5A2 is known to mediate stromal-epithelial interactions that influence prostate growth, the relationship between baseline SRD5A2 expression and prostate volume remains unclear. In this study, we analyzed SRD5A2 expression in human prostate tissues from the Medical Therapy of Prostatic Symptoms (MTOPS) trial and institutional biorepository cohorts. Quantitative assessments were performed and correlations were evaluated between expression level of SRD5A2, WNT5A, prostate volume, and tissue signaling profiles. SRD5A2 expression was significantly associated with total prostate and transition zone volume in both human cohorts. Stromal-specific WNT5A expression showed a strong positive correlation with SRD5A2, while neither serum nor tissue dihydrotestosterone levels correlated with SRD5A2 expression. In *Srd5a2-*null mice, *Wnt5a* expression in the prostate stroma was dependent on *Srd5a2* and showed region-specific regulation. Mechanistically, SRD5A2 overexpression in human prostate stromal cells upregulated WNT5A and Lymphoid Enhancer-Binding Factor 1 (LEF1), activated insulin-like growth factor 1 (IGF1) signaling, increased proliferation, and reduced apoptosis. Conditioned media from these cells enhanced epithelial proliferation through paracrine IGF1 activity, independent of epithelial WNT signaling. This study provides the first evidence that SRD5A2 promotes prostate growth through a stromal WNT5A-LEF1-IGF1 paracrine signaling axis, functioning independently of androgen levels. These findings suggest a novel therapeutic mechanism relevant for BPH patients with resistance to conventional 5α-reductase inhibitor therapy.

## Introduction

Benign prostatic hyperplasia (BPH) is a highly prevalent condition among aging men and a leading cause of lower urinary tract symptoms (LUTS). One of the primary drivers of prostate growth in adulthood is androgen signaling, mediated by the binding of androgens to the androgen receptor (AR), which activates downstream transcriptional programs that promote cellular proliferation and differentiation (1). The steroid 5α-reductase type 2 (SRD5A2), which is highly expressed in prostate tissue, catalyzes the conversion of testosterone into dihydrotestosterone (DHT), a more potent androgen that stimulates both epithelial and stromal proliferation (2). Elevated SRD5A2 activity has been associated with increased intraprostatic DHT levels and larger prostate volumes (3, 4).

Clinically, 5α-reductase inhibitors (5ARIs) such as finasteride and dutasteride are commonly prescribed to reduce prostate volume by blocking DHT synthesis and dampening androgen-driven prostate growth (5). However, finasteride reduces the progression of LUTS by only approximately 34%, suggesting that androgen-independent mechanisms may also contribute to prostate enlargement and therapeutic resistance (6).

Among the key pathways implicated in tissue growth regulation is the Wingless-related integration site (WNT) signaling pathway, which controls epithelial proliferation, stem cell maintenance, and differentiation in multiple organs (7, 8). In the prostate, canonical WNT signaling promotes basal stem cell proliferation via activation of the β-catenin-<u>T</u>-<u>c</u>ell factor**/**lymphoid <u>e</u>nhancer-binding factor (TCF/LEF) transcription complex, while non-canonical WNT signaling, particularly through WNT5A, facilitates stromal-epithelial interactions via calcium-dependent signaling cascades (9, 10). WNT5A, secreted predominantly by prostate stromal cells, has been shown to promote epithelial proliferation through paracrine mechanisms, independent of classical AR activation (10, 11). Importantly, WNT5A signaling becomes upregulated in conditions of androgen deprivation, further supporting its role in maintaining prostate growth under low-androgen states, such as during 5ARI therapy (12). In response to reduced androgen levels, stromal cells increase WNT5A secretion, supporting continued epithelial proliferation (12). Given that a subset of patients with BPH do not respond to androgen-targeting therapies, WNT5A and its downstream signaling components may provide an alternative pathway that promotes prostate growth independently of DHT levels.

Despite advances in our understanding of SRD5A2 and WNT signaling individually, the molecular relationship between these pathways and their combined contribution to prostate growth remains poorly defined. Specifically, it is unclear whether WNT5A acts downstream of SRD5A2 or constitutes an independent regulatory mechanism. While SRD5A2’s role in DHT synthesis is well established, emerging data suggest that prostate enlargement may also be driven by alternative stromal-derived signals, such as WNT5A, that function independently of androgen levels (11, 13).

In this study, we investigate the interplay between SRD5A2 and WNT signaling, with a particular focus on WNT5A as a potential mediator of SRD5A2-driven prostate growth. We hypothesize that WNT5A serves as a key DHT-independent effector linking SRD5A2 expression to prostate volume expansion. Through the integration of clinical datasets, preclinical models, and molecular assays, we aim to elucidate a novel stromal signaling axis that contributes to BPH pathogenesis and may offer a therapeutic target for patients unresponsive to conventional androgen-deprivation therapies.

## Results

### The expression of SRD5A2 is correlated with prostate volume

Our previous work demonstrated an inverse correlation between SRD5A2 expression and the clinical efficacy of 5ARIs in treating BPH (14). To address if baseline SRD5A2 expression in prostatic tissue is associated with prostate volume, we first examined 61 prostate biopsy samples from the Medical Therapy of Prostatic Symptoms (MTOPS) study cohort (Table 1) (6). We observed a statistically significant positive correlation between SRD5A2 expression and total prostate volume (Figure 1A, B). Notably, SRD5A2 expression was also significantly correlated with the transition zone volume, but not with the peripheral zone volume (Figure 1C, D), which aligns with our earlier report (15). Interestingly, despite SRD5A2’s role in DHT synthesis, we did not observe a significant correlation between clinically recorded DHT levels and SRD5A2 expression in the prostate (Figure 1E).

**Figure 1.**
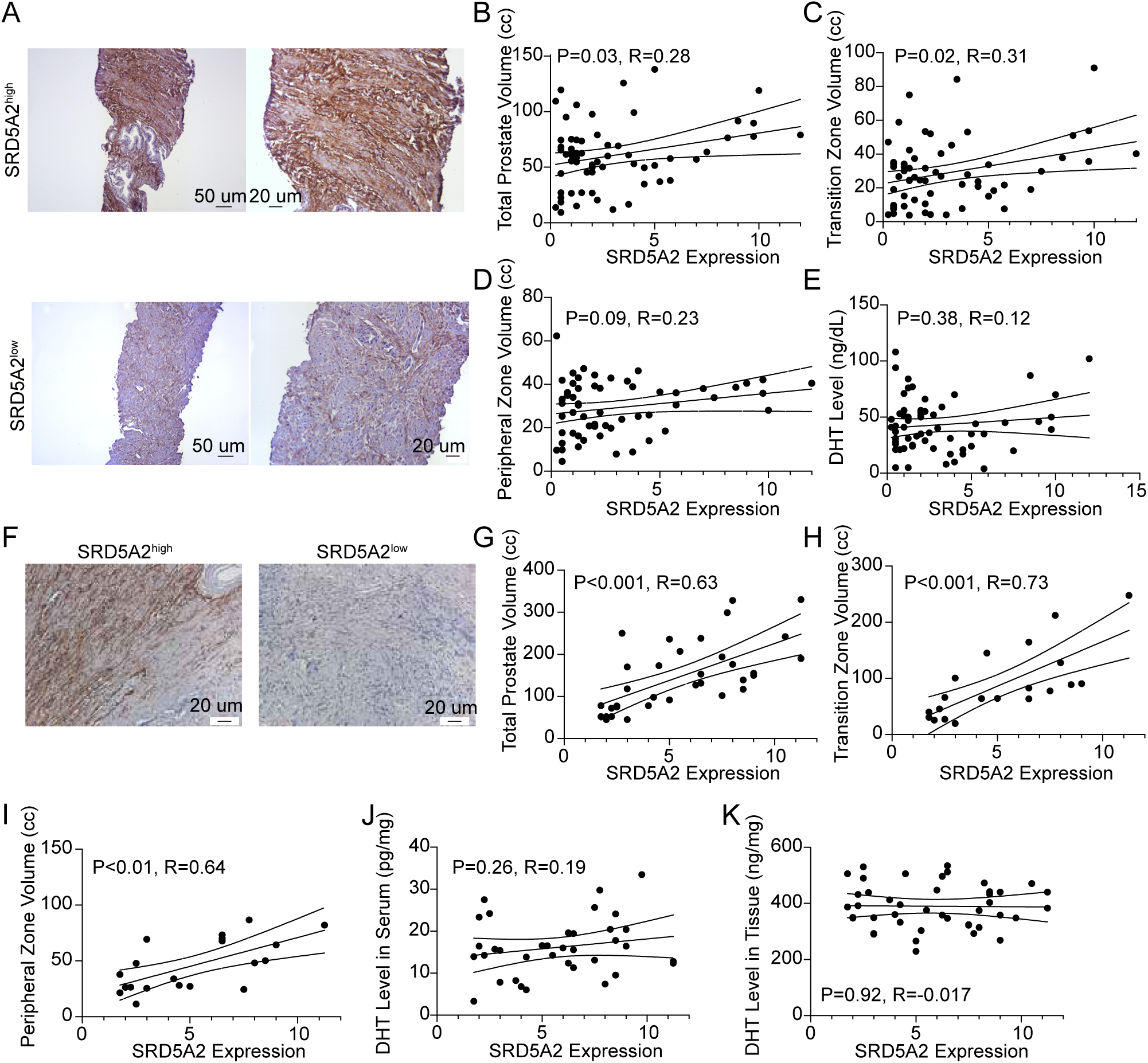
Prostate size is correlated with SRD5A2 expression in clinical samples. (**A**) Representative image of high versus low SRD5A2 expression in the MTOPS cohort (Left: magnification 10×, scale bar: 50 μm; Right: Left: magnification 20×, scale bar: 20 μm). (**B-E**) Correlation plot of SRD5A2 protein expression with total prostate volume (**B**), transition zone volume (**C**), peripheral zone volume (**D**), and serum DHT level (**E**) in the MTOPS cohort (n=61). (**F**) Representative image of high versus low SRD5A2 expression in the BIDMC biorepository cohort (magnification 20×, scale bar: 20 μm). (**G-K**) Correlation plot of SRD5A2 protein expression with total prostate volume (**G**, n=35), transition zone volume (**H**, n=20), peripheral zone volume (**I**, n=20), serum DHT level (**J**, n=23) and tissue DHT level (**K**, n = 43) in the BIDMC biorepository cohort. SRD5A2 expression was measured by immuoreactive score quantified from IHC. Serum and tissue DHT level of BIDMC biorepository samples were measured by ELISA kit. Correlation analysis was performed using Pearson correlation.

To validate our findings, we analyzed an independent cohort of 49 BPH prostate specimens from BIDMC institutional biorepository (Table 2). Consistent with the MTOPS data, SRD5A2 expression in these samples was strongly correlated with total prostate volume and transition zone volume (Figure 1 F–H). Peripheral zone volume was also correlated with SRD5A2 expression in this cohort (Figure 1I). However, neither serum nor tissue DHT levels showed significant association with SRD5A2 expression (Figure 1J, K). Together, these results demonstrate that SRD5A2 protein expression is positively associated with prostate volume, particularly in the transition zone.

### SRD5A2 knockout alters stromal–epithelial crosstalk and activates muscle-related signaling pathways in the mouse prostate

To understand the molecular mechanism of SRD5A2 expression in prostate growth, we generated *Srd5a2*-null (*Srd5a2*^-/-^) mice (14). We first performed bulk RNA sequencing on wild-type (*Srd5a2*^+/+^) controls, heterozygous (*Srd5a2*^+/-^) controls, and *Srd5a2*^-/-^ mouse prostate tissues to identify putative pathways regulated by SRD5A2. Hierarchical clustering effectively distinguished the three groups based on their differential gene expression profiles (Figure 2A, Supplementary Figure 1). Gene ontology (GO) analysis identified 133 significantly upregulated and 2 downregulated genes in *Srd5a2*^-/-^ mice (Supplementary Figure 2A, B). Notably, upregulated genes were highly enriched in biological processes related to the muscle system. Key enriched terms include those related to muscle system processes, contractile fibers, and striated muscle development, highlighting a shift toward muscle-related cellular architecture and function (Figure 2A).

**Figure 2.**
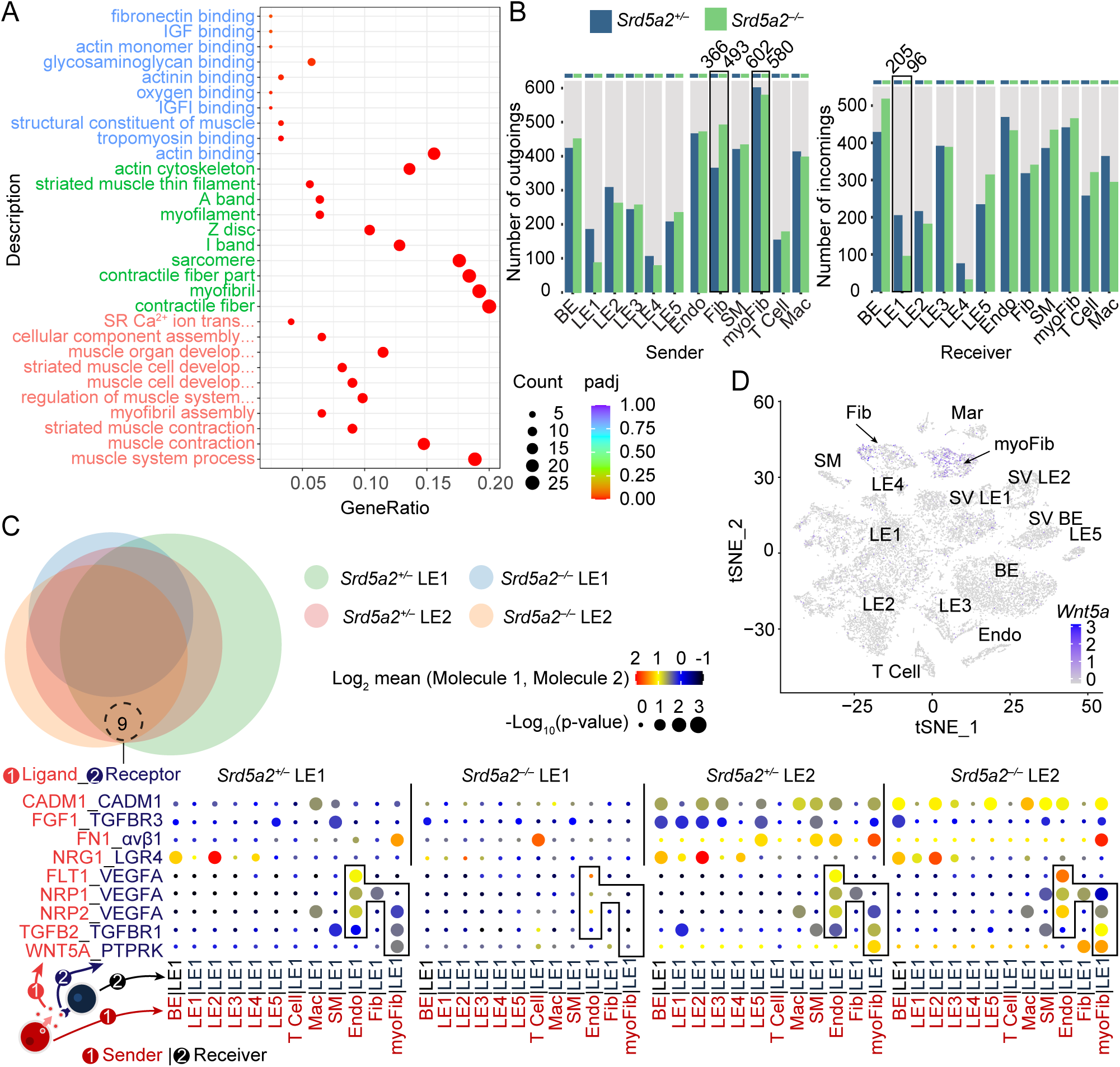
Gene and cell type profiling of *Srd5a2^-/-^* mouse prostate. (**A**) Dot plot showing the Gene Ontology (GO) enrichment analysis conducted on prostate tissue from 8-week-old mice comparing 3 control and 3 *Srd5a2*^-/-^ samples. The x-axis represents the gene ratio (the proportion of genes associated with each GO term relative to the total gene set analyzed), and the y-axis lists the GO terms grouped by their biological processes, molecular functions, or cellular components. The size of the dots indicates the number of genes (Count) involved in each term, and the color gradient represents the adjusted p-value (padj). Color represents the category of enrichment: blue, molecular function; green, cellular component; red, biological process. (**B**) The bar graphs showing the number of outgoing (secreted/paracrine/autocrine) signals and the number of incoming (ligand binding/receptor activation) signals among the cell clusters. (**C**) The CellPhone-DB analysis of the ligand-receptor interactions in different mouse prostate cell populations (n=9). Each dot represents the interaction under a specific condition, with the size indicating the statistical significance (−log_10_(p-value)) and the color representing the log_2_ mean expression level of the molecules. P-values are derived from one-sided permutation tests (significant p < 0.05). Larger and darker-colored dots signify stronger and more significant interactions. (**D**) The t-SNE (t-distributed Stochastic Neighbor Embedding) dimensionality reduction plot showing the expression localization of *Wnt5a* in different cell types of mouse prostate tissue. The expression levels is shown in color gradient within cells.

To further delineate how *Srd5a2* deletion affects cell-cell communication, we performed single cell RNA sequencing (scRNA-Seq) on mouse prostate tissues. It is well-established that paracrine signaling between stroma and epithelium governs prostate development and regeneration (10). Therefore, we focused on the interactions between luminal epithelial cells (LE) and stromal cell populations. With CellPhoneDB2 mapping the intercellular communication networks across genotypes (16), stromal cells emerged as major signal senders, and particularly, myofibroblasts showed the highest outbound signaling in both *Srd5a2*^-/-^ mice and *Srd5a2*^+/-^ controls. Notably, the total weighted ligand activity of fibroblasts increased substantially in *Srd5a2*^-/-^ animals (from 366 to 493), suggesting enhanced fibroblast-mediated signaling upon *Srd5a2* deletion (Figure 2B).

On the receiving end, *Srd5a2* knockout led to a striking reduction in receptor expression in LE1 epithelial cells (from 205 to 96), whereas LE2 cells maintained a stable number of receptors (Figure 2B). When examining stromal-derived ligands that may influence LE1 and LE2 behaviors, we identified nine significant pairs with potential regulatory roles (Figure 2C). Among these interactions, WNT5A/ protein tyrosine phosphatase receptor type K (PTPRK) stood out as candidate with differential activity across luminal subtypes (Figure 2C). Pathway analysis further confirmed activation of WNT signaling in LE2 cells upon *Srd5a2* loss (14). As a particular interest, WNT5A is a non-canonical WNT ligand that known to modulate prostate growth and epithelial behavior (10, 16–20). *Wnt5a* was predominantly expressed in stromal compartments, particularly myoFib and Fib populations (Figure 2D) (10). In addition, several transcription factors including *Tcf7l1*, a downstream transcription factor of the WNT canonical pathway, were also observed to be enriched in LE2 cells (Supplementary Figure 3A, B). These spatially restricted expression patterns suggest a targeted influence on nearby luminal cells.

### Spatial expression and regulation of WNT5A in the mouse and human prostate tissue

Next we performed RNA-FISH to assess *Wnt5a* spatial expression. Representative images reveal that *Wnt5a* transcripts in red puncta are predominantly localized in the stromal compartment, with minimal signal observed in epithelial cells (Figure 3A). There was a significant reduction of *Wnt5a* expression in the periurethral region (PrU) of *Srd5a2*^−/−^ prostates compared to both *Srd5a2*^+/+^ and *Srd5a2*^+/-^ controls (Figure 3B). While the dorsal and lateral lobes (DLP) exhibited no statistically significant differences between genotypes (Figure 3C), anterior lobes (AP) showed significant increase of *Wnt5a* signal in *Srd5a2*^−/−^ prostates (Figure 3D). Dissecting mouse prostate morphology and comparison with human prostate have confirmed that the AP of mouse prostate is more similar to the transition zone of human prostate, where BPH mostly occurs and enriches epithelial cells, while PrU and DLP are higher in stromal cell populations compared to AP (21–23). Thus, we differentiate the spatial expression of *Wnt5a* between AP and non-AP, corresponding to epithelium and stroma region of mouse prostate. *Srd5a2* knockout downregulated the mRNA level of *Wnt5a* in whole prostate (Figure 3G, H), and the difference in spatial expression of *Wnt5a* was further confirmed by qPCR in AP and non-AP lobes (Figure 3E, F). These results suggest that *Wnt5a* expression is dependent on Srd5a2 function, and exhibits regionally distinct regulation across different prostate compartments, particularly in the periurethral (proximal) region (10, 22).

**Figure 3.**
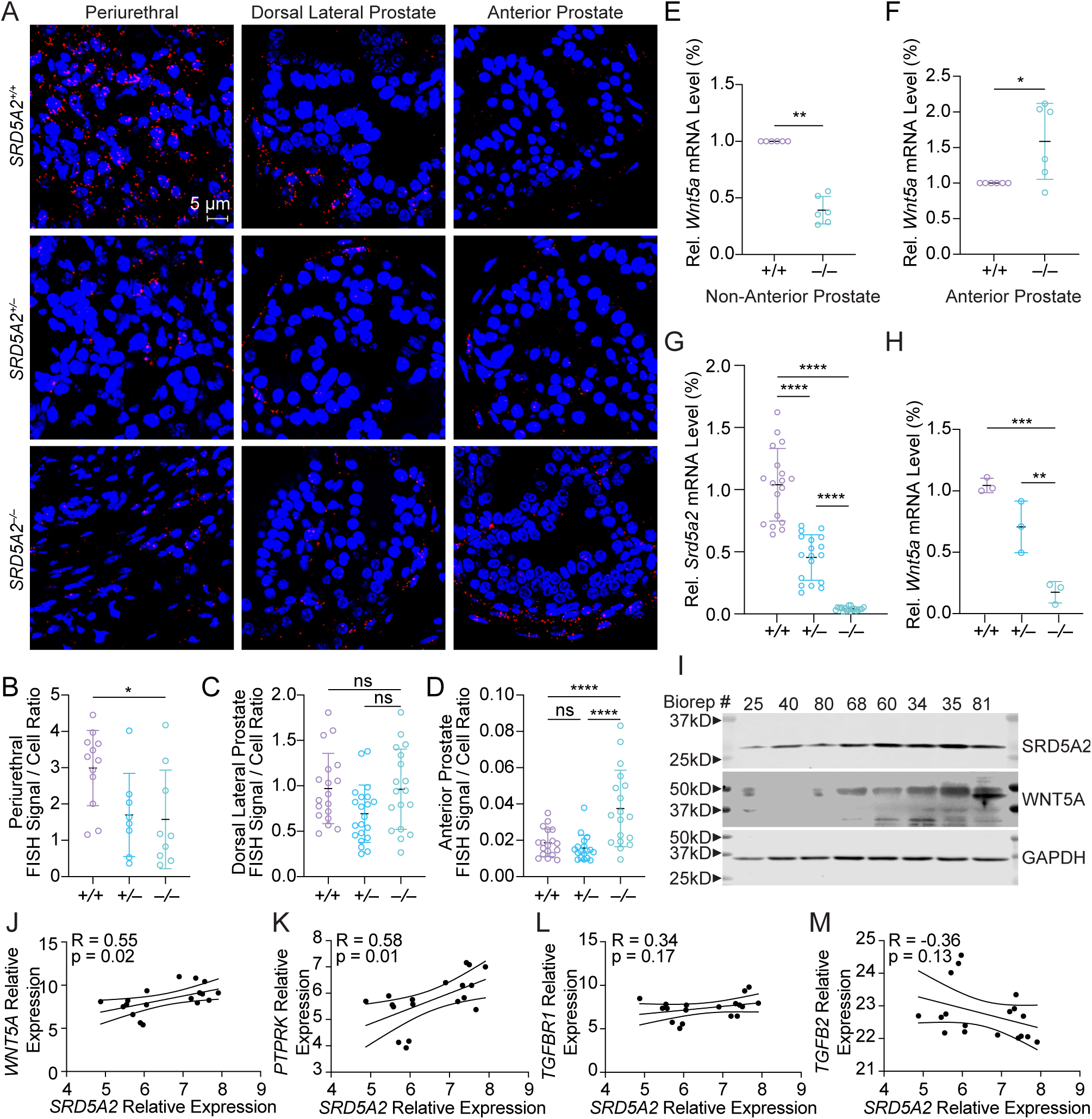
SRD5A2 level positively correlates with the expression of WNT5A in mouse and human prostate. (**A**) The representative RNA-FISH images showing *Wnt5a* (white), LE2-specific marker *Esr1* (red), and *Pvt1* (green) expression in the periurethral, dorsal lateral and anterior lobes of control and *Srd5a2*^-/-^ mice (magnification 100×, scale bar: 5 μm). (**B-D**) Quantification of *Wnt5a* RNA-FISH signal per cell in each lobe shown in (A). (**E-F**) qPCR analysis of *Wnt5a* in non-anterior and anterior prostate tissue from *Srd5a2*^+/+^ and *Srd5a2*^-/-^ mice. (**G-H**) qPCR analysis of *Srd5a2* and *Wnt5a* in total prostate tissue from *Srd5a2*^+/+^, *Srd5a2*^+/-^, and *Srd5a2*^-/-^ mice. (**I**) Western blot analysis of SRD5A2, WNT5A, and GAPDH loading control in 8 institutional biorepository patient prostate samples. The number labeling of samples is retrieved from the biorepository patient record. (**J-M**) The correlation plot of *SRD5A2* mRNA expression with *WNT5A* (**J**), *PTPRK* (**K**), *TGFB2* (**L**), and *TGFBR1* (**M**) mRNA expression level in 18 patient samples. All experiments were replicated at least three times (N≥3). Data on dot plots are presented as mean ± SD. Significance in all quantifications was calculated using two-tailed Student’s t-tests. *, p ≤ 0.05; **, p ≤ 0.01; ***, p≤ 0.001; ****; p ≤ 0.0001.

Similar with that in mouse prostate tissue, the expression of SRD5A2 corresponds with WNT5A in human prostate both in protein and mRNA levels (Figure 3 I, J). Additionally, *Srd5a2* expression was also significantly correlated with the *Ptprk*, the pair of *Wnt5a* ligand found in CellPhoneDB2 (Figure 3K). In contrast, no significant correlation was observed between *Srd5a2* and *Tgfbr1* or *Tgfb2* (Figure 3 L, M), both were among the nine significant pairs with potential regulatory roles and associated with stromal-epithelial interactions (Figure 2C). Collectively, these data demonstrate that there is a regulatory relationship between SRD5A2 and WNT5A both in the human and mouse prostate.

### SRD5A2 enhances prostate stromal cell growth and survival through activation of WNT5A pathways

To investigate the regulatory effect of SRD5A2 on WNT5A expression, we overexpressed SRD5A2 in immortalized normal prostate cells (BHPrS1) and assessed WNT5A expression at both the mRNA and protein levels. qPCR and western blot and analyses confirmed robust overexpression of SRD5A2 compared to vector control (Figure 4A, B). Notably, SRD5A2 overexpression resulted in a substantial increase in both WNT5A transcript and protein levels within stromal cells (Figure 4A, B). Furthermore, ELISA quantification demonstrated significantly elevated WNT5A concentrations both intracellularly and in the culture supernatant (Figure 4C, D), indicating enhanced secretion.

**Figure 4.**
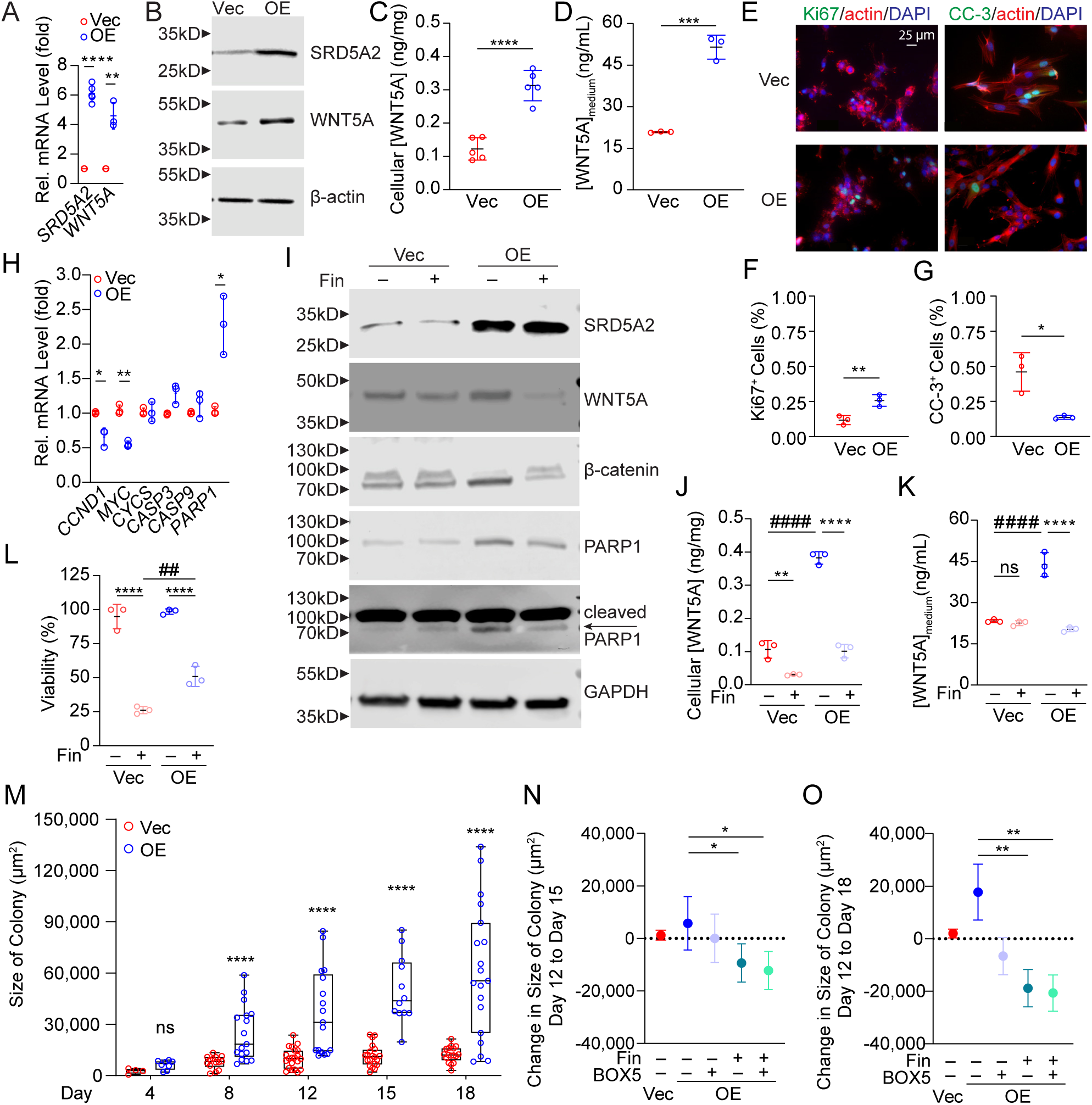
SRD5A2 regulates prostate stromal cell growth via WNT5A. (**A**) qPCR analysis of *SRD5A2*, *WNT5A* in vector control (Vec) and SRD5A2-overexpressing (OE) BHPrS1 cells. (**B**) Western blot analysis of SRD5A2, WNT5A, and actin loading control in BHPrS1 cells. (**C-D**) The level of WNT5A in BHPrS1 cell lysate and culturing medium measured by ELISA. (**E**) Representative ICC images of BHPrS1 cells showing the expression level of Ki67, and cleaved caspase 3 (CC-3). Ki67 and CC-3 protein (green) are detectable in the whole cell, Phalloidin-labeled actin filaments (red) delineate cell structure, and DAPI (blue) marks nuclear localization (magnification 40×, scale bar: 25 μm). (**F-G**) Quantification of Ki67 and CC-3 signals in (E). (**H**) qPCR analysis of proliferation and apoptosis-related markers in BHPrS1 cells. (**I**) Western blot analysis of SRD5A2, WNT5A, β-catenin, PARP1 (full and cleaved), and GAPDH loading control in BHPrS1 cells treated with DMSO or 100µM finasteride (Fin) for 72 hours. (**J-K**) The level of WNT5A in DMSO or finasteride-treated BHPrS1 cell lysate and culturing medium measured by ELISA. (**L**) Cell viability of DMSO or Fin-treated BHPrS1 cells measured by MTT assay. (**M**) Quantification of the BHPrS1 spheroid-like colony size from day 4 to day 18. (**N-O**) Quantification of BHPrS1 spheroid-like colony size change from day 12 to day 15 and from day 12 to day 18 as BHPrS1 OE cells treated with 100µM finasteride and/or 100µM BOX5 for 72 hours or 144 hours. All experiments were replicated at least three times (N≥3). Data on dot plots are presented as mean ± SD. Student’s t-test and one-way ANOVA with Tukey test were performed. ns, p > 0.05; *, p ≤ 0.05; **, p ≤ 0.01; ***, p≤ 0.001; ****; p ≤ 0.0001. #, significance in comparisons between different groups with the same meaning as *.

To analyze the effect of SRD5A2 on prostate cell growth, we performed immunofluorescence (IF) staining for the proliferation marker Ki67, and the apoptosis marker cleaved caspase-3 (CC-3). The data revealed a significant increase in Ki67 and decrease in CC-3 staining in SRD5A2 overexpression cells compared to the vector control (Figure 4E-G). Meanwhile, SRD5A2 overexpression greatly increased the expression of cleaved PARP1 (Figure 4H), an essential regulator of cell apoptosis and downstream of WNT canonical signaling particularly TCF/LEF transcription factor family (24, 25).

Next, we asked if pharmacological inhibition of SRD5A2 impacts on WNT5A and its downstream protein expression. Compared to the DMSO vehicle, SRD5A2 inhibitor finasteride suppressed the upregulation of WNT5A, β-catenin, and PARP1 in SRD5A2-overexpressed stromal cells (Figure 4I). ELISA quantification also revealed significantly elevated WNT5A protein levels in both stromal cells and culture medium following SRD5A2 overexpression, and markedly reduced with finasteride treatment (Figure 4J, K). From the perspective of non-canonical WNT signaling, overexpression of SRD5A2 increased calcium level, which was also abrogated with finasteride treatment (Supplementary Figure 5C). In addition, finasteride significantly reduced the viability of prostate stromal cells with normal levels of SRD5A2, whereas SRD5A2-overexpressing cells maintained significantly higher viability and showed partial resistance to finasteride-induced cytotoxicity (Figure 4L), suggesting that SRD5A2 overexpression confers a pro-survival advantage in the presence of 5ARI treatment.

To evaluate the long-term impact of SRD5A2 overexpression on prostate stromal cell proliferation in 3D culture, we performed spheroid-like colony formation assay on BHPrS1 cells. At day 4, no significant difference was observed between vector control and overexpression cells. However, from day 8 onward, SRD5A2-overexpressing cells formed significantly larger spheroid colonies than control cells, with exponential growth observed through day 18 (Figure 4M). Furthermore, treatment with BOX5, finasteride, or a combination of the two between day 12 and day 15 showed significantly reduced colony expansion (Figure 4N), and this inhibitory effect was further pronounced comparing the change between day 18 and day 12 (Figure 4O), indicating sensitivity to both androgen metabolism and WNT5A signaling disruption. These findings highlight that SRD5A2 promotes stromal spheroid growth through a WNT5A-dependent mechanism, and that targeting either 5α-reductase or WNT5A signaling can significantly impair the expansion of SRD5A2-driven stromal colonies.

### LEF1 mediates the SRD5A2-induced WNT canonical signaling pathway

To explore the molecular components of WNT signaling pathway that are regulated by SRD5A2, we performed a targeted PCR microarray analysis following SRD5A2 overexpression. Among the 84 differentially expressed genes, *LEF1, WNT5A, IGF2,* and others were significantly upregulated (Figure 5A). qPCR further confirmed that IGF1, IGF2, along with WNT5A receptors RYK, ROR1, ROR2, and LRP6 were greatly upregulated (Supplementary Figure 5E, F).

**Figure 5.**
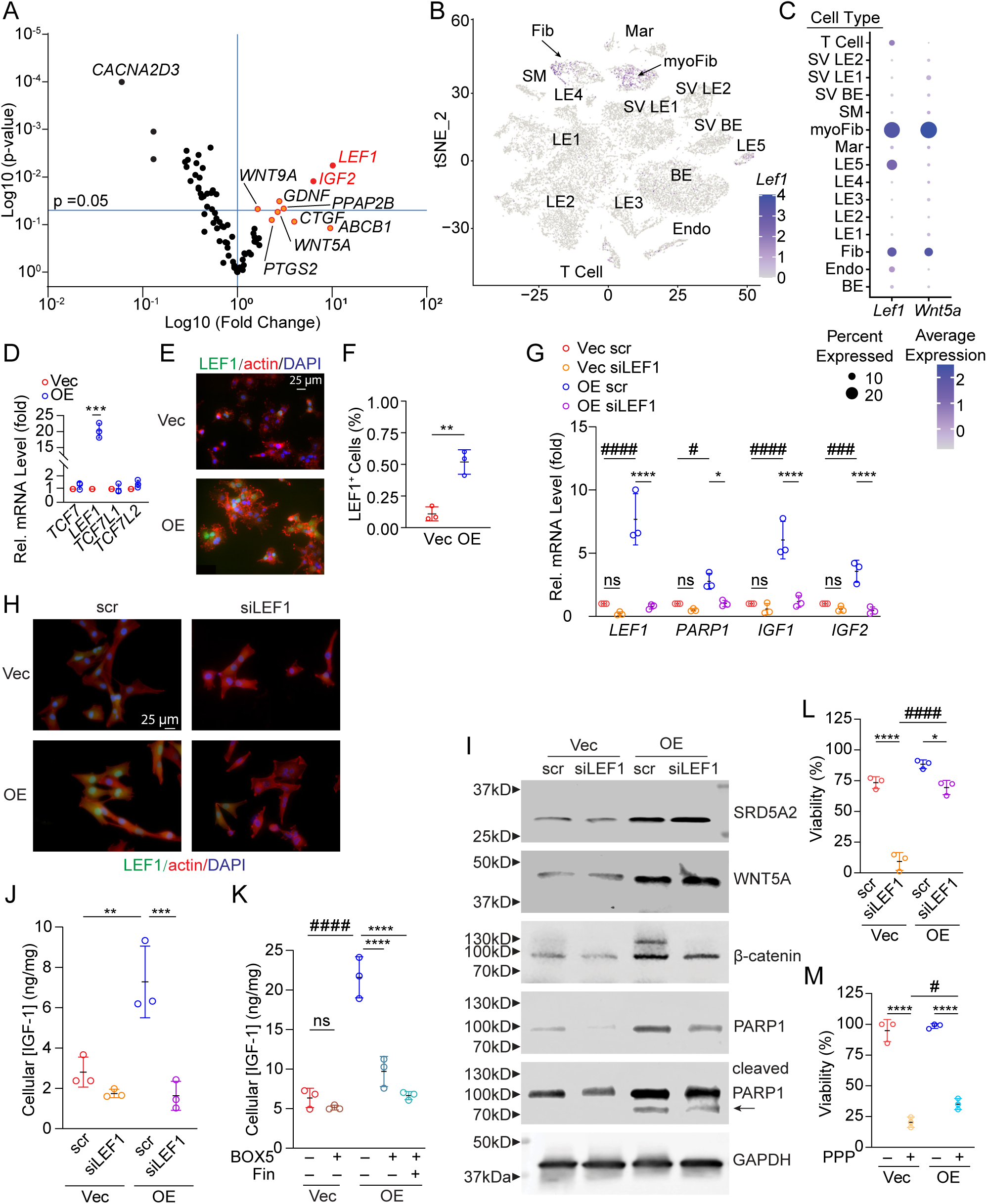
SRD5A2 upregulates the expression of LEF1 and IGF1. (**A**) Volcano plot showing differentially expressed WNT-related genes in BHPrS1 OE versus Vec cells with log_10_ (Fold Change) indicating mean gene expression levels. Each dot represents a single gene. (**B**) t-SNE dimensionality reduction plot showing the expression localization of *Lef1* in different cell types of mouse prostate tissue and its expression levels within cells. (**C**) Dot plot showing the expression of *Lef1* and *Wnt5a* in different cell types from scRNA-seq of mouse prostate tissue. The size of the dots represents the proportion of positively expressing cells, and the color indicates the average expression levels. (**D**) qPCR analysis of TCF/LEF family transcription factors in BHPrS1 cells. (**E**) Representative ICC images of BHPrS1 cells showing the expression level of LEF1. LEF1 protein (green) is detectable in the nuclei, Phalloidin-labeled actin filaments (red) delineate cell structure, and DAPI (blue) marks nuclear localization (magnification 40×, scale bar: 25 μm). (**F**) Quantification of LEF1 signals in (E). (**G**) qPCR analysis of *LEF1*, *PARP1*, *IGF1*, and *IGF2* in BHPrS1 Vec and OE cells transfected with scramble (scr) or LEF1 (siLEF1) siRNAs. (**H**) Representative ICC images showing the expression level of LEF1 in siRNA-transfected BHPrS1 cells (magnification 40×, scale bar: 25 μm). (**I**) Western blot analysis of SRD5A2, WNT5A, β-catenin, PARP1 (full and cleaved), and GAPDH loading control in siRNA-transfected BHPrS1 cells. (**J-K**) The cellular level of IGF1 in BHPrS1 cell transfected with siRNAs or treated with 100µM finasteride and/or 100µM BOX5 for 72 hours measured by human IGF1 ELISA kit. (**L-M**) Cell viability of siRNA-transfected or 72-hours Picropodophyllin (PPP)-treated BHPrS1 cells measured by MTT assay. All experiments were replicated at least three times (N≥3). Data on dot plots are presented as mean ± SD. Student’s t-test and one-way ANOVA with Tukey test were performed. ns, p > 0.05; *, p ≤ 0.05; **, p ≤ 0.01; ***, p≤ 0.001; ****; p ≤ 0.0001. #, significance in comparisons between different groups with the same meaning as *.

The *Lef1* expression within the mouse prostate was further validated by re-analyzing the scRNA-seq dataset. *Lef1* signal was shown as primary enrichment in stromal populations, particularly in myoFib and Fib (Figure 5B). Dot plot analysis further confirmed this cell type-specific expression patterns for both *Lef1* and *Wnt5a* (Figure 5C), suggesting potential functional interaction or co-regulation between *Lef1* and *Wnt5a* in these cell types. Notably, as a key transcriptional mediator of canonical WNT/β-catenin signaling (26), *LEF1* demonstrated one of the highest fold changes and statistical significance levels at both mRNA and protein level, suggesting strong activation of WNT transcriptional programs downstream of SRD5A2 (Figure 5A, D-F). The activation of canonical WNT signaling is consistent with our observations on SRD5A2-downregulated β-catenin degradation. The protein level of β-catenin is increased upon SRD5A2 overexpression and reduced by finasteride (Figure 4I).

Interestingly, other downstream transcriptional regulators of canonical WNT signaling besides LEF1, specifically *TCF7*, *TCF7L1* and *TCF7L2,* showed no significant change at mRNA level (Figure 5D). These findings suggest activation of the canonical WNT signaling pathway by SRD5A2 is specific to the transcription factor LEF1 rather than the β-catenin-bound TCF/LEF family in general.

To determine whether LEF1 mediates the downstream effects of SRD5A2 overexpression, we transfected BHPrS1 stromal cells with either scrambled (scr) or LEF1-targeting siRNA (siLEF1) and performed qPCR, IF, and Western blot analysis subsequently. LEF1 is markedly reduced after siLEF1 transfection in SRD5A2 overexpression cells, confirming efficient knockdown at mRNA and protein level (Figure 5G, H). Western Blot results showed that SRD5A2 and WNT5A protein levels remained high in SRD5A2 overexpression group and were unaffected by siLEF1 (Figure 5I), and silencing LEF1 reduced SRD5A2-induced upregulation of PARP1 expression at both mRNA and protein levels in both vector and SRD5A2-overexpression groups compared corresponding cells transfected with scramble siRNA control (Figure 5G, I). LEF1 knockdown significantly reduced cell viability in both cells, although to a lesser extent in SRD5A2 overexpression cells, indicating that LEF1 partially mediates SRD5A2-induced resistance to cell death (Figure 5L).

### LEF1 regulates IGF1 expression in prostate stromal cells through a WNT5A-dependent pathway

Given that the IGF1 promoter contains a WNT response element and that IGF1 has been reported as a direct WNT target gene (27), we next investigated whether IGF1 expression is regulated downstream of SRD5A2 via the WNT5A-LEF1 axis. Both mRNA and protein levels of IGF1 were significantly elevated in SRD5A2-overexpressing stromal cells, and markedly reduced following LEF1 knockdown (Figure 5G, J). Furthermore, pharmacologic inhibition using either Finasteride or BOX5, a WNT5A antagonist, significantly suppressed IGF1 expression (Figure 5K), supporting the notion that SRD5A2 enhances IGF1 transcription in a WNT5A/LEF1-dependent manner.

Functionally, treatment with Picropodophyllin (PPP), a selective IGF1 receptor inhibitor, led to a significant reduction in cell viability in both vector control and SRD5A2-overexpressing cells (Figure 5M). Together, these findings demonstrate that LEF1 and IGF1 signaling are essential for SRD5A2-mediated survival in prostate stromal cells, and that targeting these pathways may overcome stromal cell–mediated resistance mechanisms.

### SRD5A2-overexpressing stromal cells promote epithelial proliferation in an IGF1 paracrine manner

To evaluate the SRD5A2 induced ligand-receptor interaction change between stromal cells and adjacent epithelial cells, we co-cultured BPH1 prostate epithelial cells with BHPrS1 stromal cells and analyzed epithelial expression profiles by qPCR and Western blot. Co-culturing BPH1 cells with SRD5A2 overexpression BHPrS1 cells resulted in a slight reduction in mRNA level of *WNT5A* and *LEF1* in BPH1 cells (Figure 6A, Supplementary Figure 6A), suggesting feedback regulation of epithelial WNT activity. Unlike stromal cells, the mRNA level of WNT receptors in epithelial cells largely remained unchanged (Supplementary Figure 6B). Notably, SRD5A2-overexpressing stromal cells induced a strikingly increase in *IGF1* mRNA level in BPH1 cells (Figure 6A). The change of IGF1 expression at protein level was further confirmed by ELISA (Figure 6B). The results suggest that SRD5A2-overexpressing stromal cells might modulate epithelial growth through enhancing IGF1 receptor activation.

**Figure 6.**
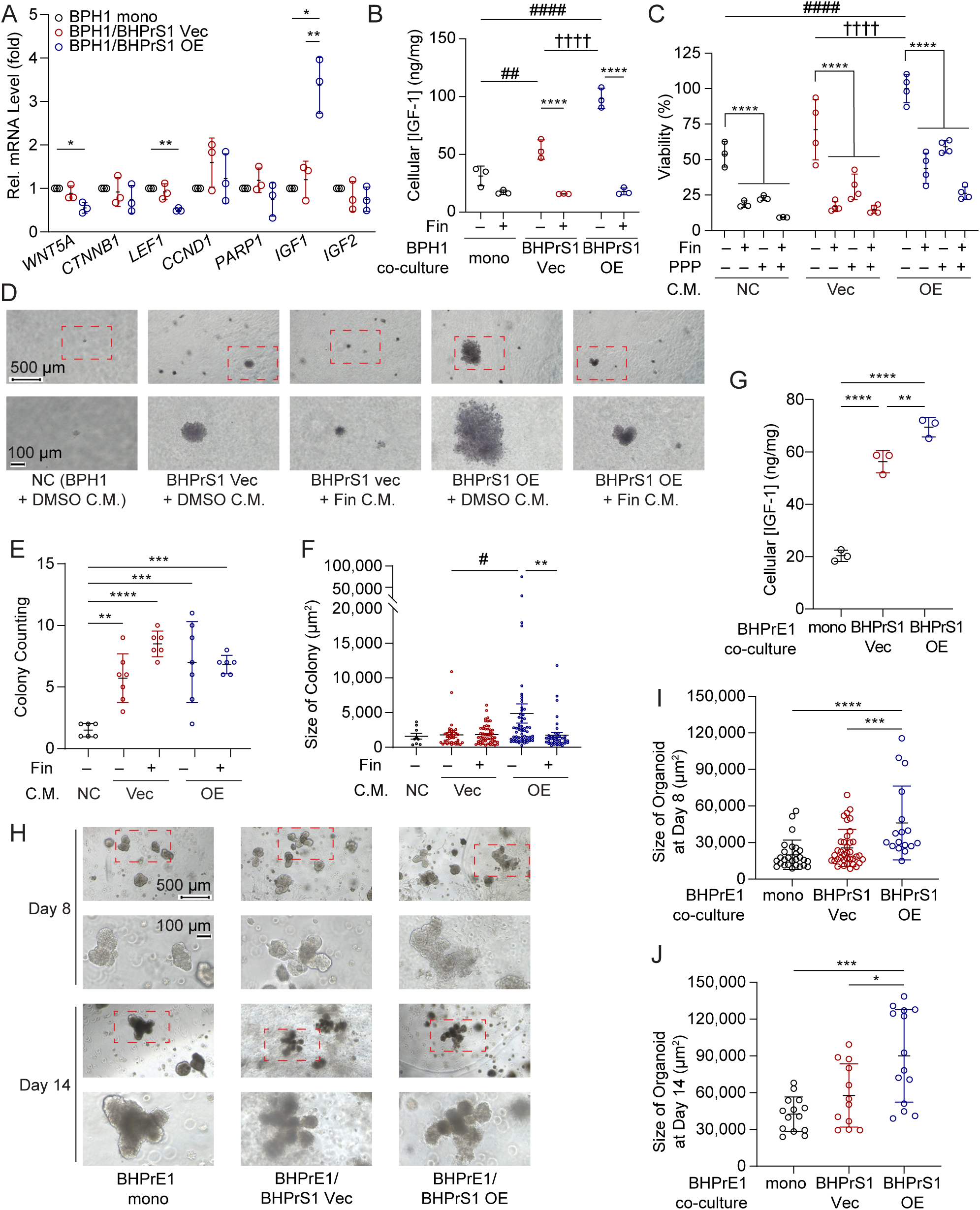
SRD5A2-overexpression stromal cells increased epithelial cells IGF1 expression and cell proliferation. (**A**) qPCR analysis of *WNT5A*, β*-catenin*, *LEF1*, *cyclin D1*, *PARP1*, and WNT-related growth factors in BPH1 cells co-cultured with BPH1 cells, BHPrS1 control cells, or BHPrS1 SRD5A2 overexpressing cells. (**B**) The cellular level of IGF1 in BPH1 cells co-cultured with BPH1 cells, BHPrS1 control cells, or BHPrS1 SRD5A2 overexpressing cells measured by human IGF1 ELISA kit. (**C**) Cell viability of BPH1 cells cultured in control or drug-treated BHPrS1-conditioned medium for 72 hours measured by MTT assay. (**D**) Representative images taken on day 21 of BPH1 soft agar colonies in control or drug-treated BHPrS1-conditioned medium starting on day 1. (**E-F**) Quantification of the colony numbers (**E**) and colony size (**F**) of BPH1 cells shown in (D). (**G**) The cellular level of IGF1 in BHPrE1 cells co-cultured with BHPrE1 cells, BHPrS1 control cells, or SRD5A2 overexpressing BHPrS1 cells for 72 hours measured by human IGF1 ELISA kit. (**H**) Representative images taken on day 8 and day 14 of BHPrE1 organoid co-cultured with BHPrS1 cells. (**I-J**) Quantification of the size of BHPrE1 organoids on day 8 (**I**) and day 14 (**J**) shown in (H). All experiments were replicated at least three times (N≥3). Data on dot plots are presented as mean ± SD. One-way ANOVA with Tukey test were performed to calculate the significance. ns, p > 0.05; *, p ≤ 0.05; **, p ≤ 0.01; ***, p≤ 0.001; ****; p ≤ 0.0001. # and †, significance in comparisons between different groups with the same meaning as *.

To further assess the effect of SRD5A2-overexpressing stromal cells on epithelial proliferation, we treated BPH1 cells with conditioned media (CM) from both vector control and SRD5A2-overexpressing BHPrS1 cells. BPH1 cell viability increased with CM from both groups, but CM from SRD5A2-overexpressing stromal cells induced a significantly greater increase (Figure 6C). Treatment with either finasteride or PPP significantly reduced the CM’s viability-promoting effect on BPH1 cells, with the strongest inhibition observed upon the combined treatment (Figure 6C). CM from LEF1 siRNA transfected stromal cells had much lower to almost no epithelial viability boosting effect compared to CM from the scramble control cells, further validating that the paracrine effect of stromal cells on epithelial viability is LEF1-dependent (Supplementary Figure 6C). Additionally, soft agar assays showed that CM significantly increased both the number and size of BPH1 colonies compared to regular medium (Figure 6D-F).

CM from SRD5A2-overexpressing stromal cells induced the greatest colony growth, which was significantly reduced by finasteride (Figure 6E-F). These findings suggest that stromal SRD5A2 promotes epithelial expansion via soluble factors, such as IGF1, in a finasteride-sensitive manner.

To validate the functional impact of stromal SRD5A2 on epithelial morphogenesis, organoid-forming BHPrE1 cells were co-cultured with BHPrS1 stromal cells in a 3D matrix. Consistent with BPH1 results, co-culture with SRD5A2-overexpressing stromal cells increased IGF1 expression (Figure 6G), and led to the formation of larger, more complex organoids by day 8, with even more pronounced growth and branching by day 14 (Figure 6H). Quantification confirmed a significant increase in organoid size at both time points (Figure 6I, J). These results support a role for stromal SRD5A2 in promoting epithelial proliferation and prostate tissue remodeling through paracrine signaling.

### WNT5A expression correlates with SRD5A2 and total prostate volume in human prostate tissues

To evaluate the clinical relevance of WNT5A in the human prostate, we first analyzed biopsy samples from the MTOPS clinical trial cohort. IHC staining showed stromal-enriched expression of WNT5A with distinct intensity differences across samples (Figure 7A). Quantitative analysis revealed a strong positive correlation between WNT5A and SRD5A2 expression (P < 0.001, R = 0.62; Figure 7B), suggesting potential co-regulation of these genes within the prostate stroma. To assess whether WNT5A expression was associated with total prostate volume, we observed a marginal significant correlation (P = 0.02, R=0.39; Figure 7C), indicating that WNT5A levels may not directly reflect prostate size.

**Figure 7.**
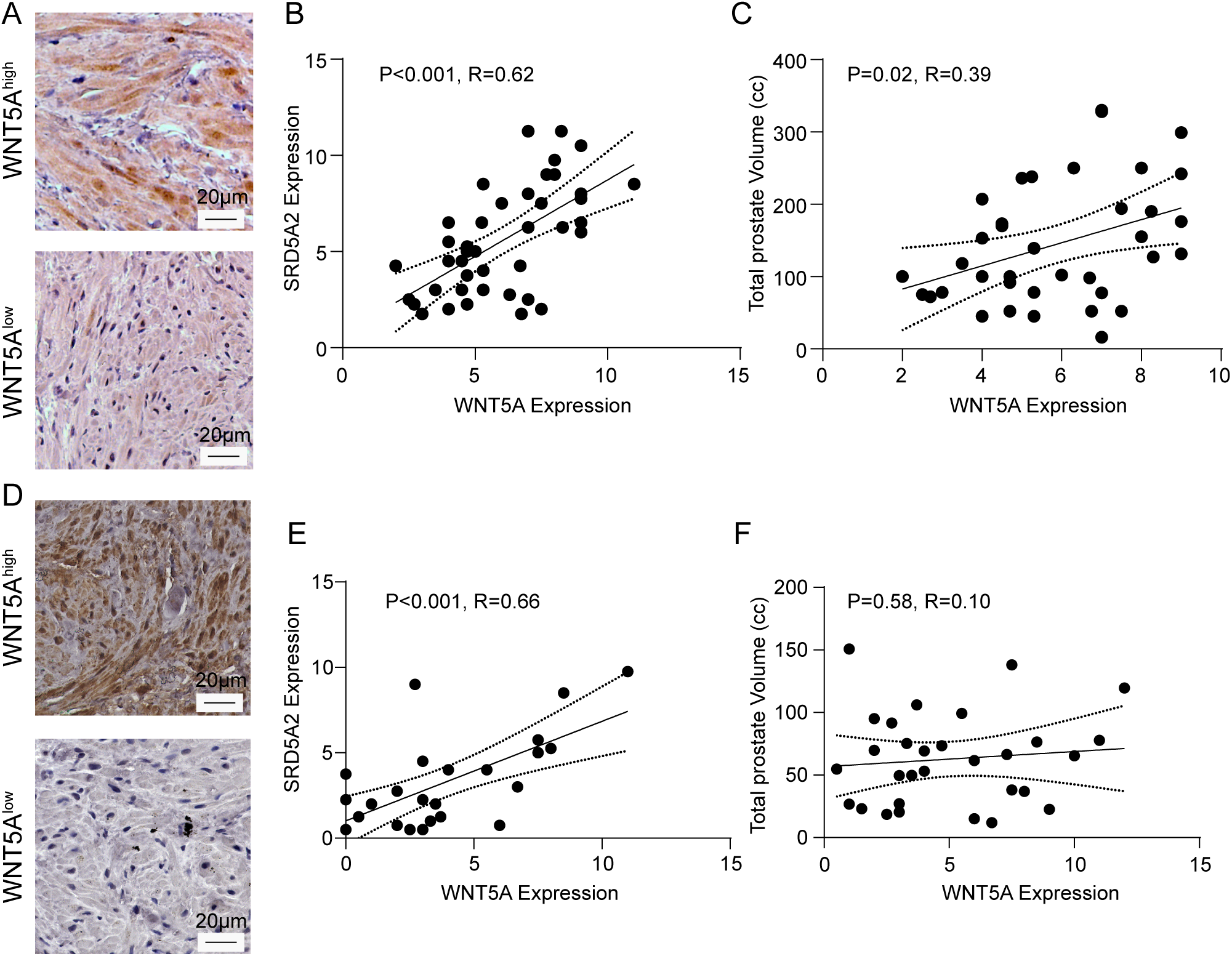
Prostate size and SRD5A2 expression level are correlated with WNT5A expression in clinical samples. (**A**) Representative image of high versus low WNT5A expression in the BIDMC biorepository cohort (magnification 20×, scale bar: 20 μm). (**B-C**) Correlation plot of WNT5A expression with SRD5A2 expression (**B**, n=41) and total prostate volume (**C**, n=37) in the BIDMC biorepository cohort. (**D**) Representative image of high versus low WNT5A expression in the MTOPS cohort (magnification 20×, scale bar: 20 μm). (**E-F**) Correlation plot of WNT5A expression with SRD5A2 expression (**E**, n=25) and total prostate volume (**F**, n=25) in the MTOPS cohort. WNT5A and SRD5A2 expressions were measured by immunoreactive score (IRS) quantified from IHC. Correlation analysis was performed using Pearson correlation.

To validate these findings, we examined an independent biorepository cohort. Similar to the MTOPS cohort, WNT5A staining remained stromal-specific (Figure 7D) and was strongly correlated with SRD5A2 expression (P < 0.001, R = 0.66; Figure 7E). Again, there was no significant correlation between WNT5A expression and total prostate volume (P = 0.58, R = 0.10; Figure 7B), reinforcing the conclusion that WNT5A is not a determinant of glandular size. Together, these results demonstrate that WNT5A and SRD5A2 are tightly co-expressed in the prostate stroma, but WNT5A levels do not correlate with prostate volume, suggesting its role is more likely related to tissue signaling than to structural enlargement.

## Discussion

In this study, we present important new insights regarding the role of SRD5A2 in BPH. We observed a significant positive correlation between SRD5A2 protein expression and prostate volume, notably within the transition zone (Figure 1), where most hyperplastic changes occur. This finding aligns well with known clinical features of BPH, where enlargement of the prostate transition zone primarily drives LUTS characteristic of BPH (28, 29). Although SRD5A2 mediates the conversion of testosterone into DHT, a potent androgen responsible for prostate growth, a direct relationship between enzyme expression and prostate volume appears biologically plausible. Our findings in two independent cohorts suggest that SRD5A2 expression might serve as an important biomarker for prostate growth and potentially predict responsiveness to 5ARI therapy.

Interestingly and somewhat unexpectedly, our analyses revealed no significant correlation between SRD5A2 and intraprostatic or serum DHT concentrations (Figure 1E, J, K). This is contrary to the traditional paradigm that directly links SRD5A2 expression to elevated local DHT levels within prostate tissue (3, 30). This result could reflect complex compensatory mechanisms within the prostate microenvironment, including alternate androgen metabolic pathways, such as estrogen signaling pathway (14, 31). It is also possible that factors beyond enzymatic activity, such as enzyme kinetics, cellular localization, or variations in androgen receptor sensitivity and signaling pathways, might affect intraprostatic androgen levels (32, 33). Therefore, our study highlights SRD5A2 as a potential determinant of prostate size isolated from traditional androgen-dependent mechanisms. Further studies exploring these mechanisms are warranted to better elucidate how SRD5A2 contributes to prostate tissue growth independent of local androgen concentration.

Our study identifies a novel regulatory pathway involving SRD5A2, WNT5A, LEF1, and IGF1 in the stromal compartment of BPH (Supplementary Figure 6D). We demonstrate that SRD5A2 overexpression significantly increases the production and secretion of WNT5A, activating downstream signaling via LEF1, PARP1 and IGF1 to promote stromal cell proliferation and resistance to apoptosis (Figure 5, 6). LEF1 is known to drive transcriptional programs critical for cell proliferation and survival (34). Through scRNA-seq, we localized LEF1 predominantly to prostate stromal cells, specifically Fib and myoFib. This stromal specificity corroborates previous studies suggesting stromal cells as essential mediators of prostate growth through paracrine signaling pathways (35).

The functional relevance of LEF1 was validated through siRNA-mediated knockdown, which significantly reduced SRD5A2-induced PARP1 and IGF1 expression and attenuated stromal cell viability (Figure 5). PARP1 is involved in repairing DNA damage and a well-known therapeutic target for inducing apoptosis of cancer cells to reduce prostate cancer growth and invasion (25).

IGF1 is a well-characterized growth-promoting and anti-apoptotic factor involved in prostate biology and BPH pathology (36–38). Notably, pharmacological inhibition of IGF1 signaling via PPP, an IGF1 receptor inhibitor, significantly decreased viability of SRD5A2-overexpressing stromal cells, further underscoring the critical role of IGF1 in stromal cell proliferation and survival mediated by SRD5A2-WNT5A-LEF1 signaling.

Importantly, the observed partial resistance to finasteride in SRD5A2-overexpressing stromal cells suggests that targeting the downstream signaling components, such as WNT5A or IGF1, might result in a synergic effect and offer additional therapeutic benefits. In conclusion, our study presents compelling evidence for an SRD5A2-WNT5A-LEF1-IGF1 signaling axis within prostate stromal cells that enhances cell proliferation and survival independently of classical androgen receptor signaling.

While our findings suggest that WNT5A acts primarily as a signaling mediator rather than a direct driver of stromal or glandular growth, further validation of AR-independent role of WNT5A and IGF1 and their importance in hyperplasia development is needed, especially from *in vivo* perspective. As one of essential stromal-derived factors, WNT5A and IGF1 are known profoundly influence epithelial biology through paracrine mechanisms prostate (11, 39–41). A recent study further reported correlations between androgen, IGF1, and WNT/β-catenin signaling, as well as a dynamic regulatory loop of IGF1-WNT initiated by the AR through prostatic oncogenesis development in human prostate cancer samples (42), while other studies found IGF1 and WNT5a enhanced both androgen-dependent and androgen-independent prostatic development in both cancer and noncancerous cells (11, 43, 44). Thus, the effects of paracrine WNT5A and IGF1 on organ size and underlying mechanism are complex and context-dependent. Therefore, future *in vivo* studies are necessary to directly assess the impact of WNT5A and the WNT5A-LEF1-IGF1 signaling axis on prostate growth and development.

Currently, 5ARIs represent mainstay medical treatments for BPH, yet patient responses remain highly variable (45, 46). 5ARIs also showed synergic effects with alpha-blockers or PDE5 inhibitors that address the smooth muscle contraction, indicating a good potential for combination therapy with other drugs (47–49). Our earlier work demonstrated an inverse correlation between SRD5A2 expression and clinical response to 5ARIs (14). The current findings extend this understanding, suggesting that SRD5A2 expression, independent of local DHT concentrations, may inform treatment decisions or the development of complementary therapeutic strategies aimed at other stromal-epithelial signaling pathways. In conclusion, our findings highlight the critical stromal co-expression of WNT5A and SRD5A2, uncover an essential paracrine role for stromal SRD5A2 in regulating prostate epithelial proliferation and morphogenesis via the IGF1 signaling pathway. These findings highlight a critical stromal-epithelial axis in prostate tissue remodeling and provide new insights for more precise therapeutic interventions in BPH management.

## Materials and Methods

### Clinical Samples

De-identified human prostate biopsy samples (n = 61) collected during the MTOPS were obtained from the National Institutes of Health/National Institute of Diabetes and Digestive and Kidney Diseases (NIH/NIDDK) Central Repository under an approved X01 application. Relevant clinical data from the MTOPS trial, including total prostate volume, transition zone volume, and serum DHT levels, were obtained and utilized for correlative analysis. Surgical prostate specimens were obtained from the Beth Israel Deaconess Medical Center Prostate Biorepository (n = 49). Relevant clinical data, including total prostate volume and transition zone volume, derived from prostate MRI, CT scans of the abdomen and pelvis, and genitourinary ultrasounds prior to the surgical date, were collected and utilized for correlative analysis.

### Mouse Model

Generation of Srd5a2-null (*Srd5a2^-/-^*) and heterozygous (*Srd5a2^+/–^*) mice model was previously described (14). Mice were housed in standard polycarbonate cages with free access to normal food and water. We only studied biological males since the prostate is the organ specific to biological male animals. All animal work described in this study was in adherence to US NIH Guide for the Care and Use of Laboratory Animals and approved by the Beth Israel Deaconess Medical Center Institutional Animal Care and Use Committee (protocol number 049-2022). No randomization was used

### 2D Cell Culture

Human prostate epithelial cell lines BPH-1, BHPrE1, and stromal cell line BHPrS1 were gifted by Dr. Simon Hayward of North Shore University Health System Research Institute, IL, Chicago since 2019. BHPrS1, and BPH-1 cells were maintained in RPMI1640 (Gibco, 11875119) with 10% v/v fetal bovine serum (FBS, Gibco, A5256801) and 1% Penicillin/Streptomycin (Pen/Step, Corning, 30-002-CI). Vector control and SRD5A2-overexpressing BHPrS1 stable cell lines were established by transfecting cells with 1% SRD5A2-sequence coded lentiviral particles (GeneCopoeia, A3930) and 0.1% polybrene (Millipore, TR1003). Positive cells were selected and maintained using 0.5% puromycin (Gibco, A1113802). Transfection efficiency was tested using quantitative PCR. BHPrE1 cell was cultured in DMEM/F-12 (Gibco, 11320033) with 5% FBS, 1% v/v Pen/Step, 0.4% Bovine Pituitary Extract, (Gibco, 13028014), 1% Insulin-Transferrin-Selenium (Gibco, 41400045), and 10 ng/ml Human EGF (Gibco, PHG0314). All cell lines were cultured in corresponding medium in a humidified incubator with 5% CO_2_ at 37°C. All cell lines used in this study were authenticated and intermittently evaluated in-house for mycoplasma contamination (InvivoGen, ant-mpt).

For 2D stromal-epithelial transwell co-culture, polycarbonate membrane cell culture inserts (ThermoFisher, 140660) in 6-well plates were used following manufacturer’s protocol with minor modifications. Briefly, 1.5×10^5^ cells/well of BPH-1 cells were seeded into the bottom of the 6-well plates and 1×10^5^ cells/well of BPH-1 or BHPrS1 cells were seeded inserts above. The insert and bottom of the well were each maintained with 1mL of RPMI1640 with 10% FBS. The cells in the inserts were treated with drugs or siRNAs, and the BPH-1 epithelial cells in the bottom well were collected for further analysis.

### RNA Transfection and Drug Treatment

For gene silencing with small interfering RNAs (siRNAs), BHPrS1 cells were seeded in 6-well plates and cultured for 24 hours before transfection with control (Santa Cruz, sc-37007) or LEF1 (Santa Cruz, sc-35804) siRNAs using Lipofectamine RNAiMAX Reagent (Invitrogen, 13778150) and Opti-MEM (Gibco, 51985034) according to the manufacturer’s instructions. The transfected cells were cultured for 48hours at 37°C with 5% CO_2_ before proceeding with ICC, RNA extraction, MTT assay, or sample preparation steps for Western blot.

For drug treatment and AR activation, BHPrS1 cells seeded in 6-well plates were starved for 24 hours before adding the corresponding drugs. Starved cells were then treated with 48 to 72 hours of 100 µM Finasteride (SigmaAldrich, F1293), or 100 µM BOX5 (SigmaAldrich, 681673), or 50nM PPP (ThermoFisher, J65916MA) with fresh drugs and medium changed every 24 hours. Similarly, androgen stimulation was performed by preparing the corresponding concentration of the testosterone and DHT in RPMI1640 medium with 10% charcoal stripped FBS instead of regular FBS, and adding to the cells for 72 hours treatment after starvation with fresh androgen and medium changed every 24 hours.

### Organoid and Spheroid Culture

The BHPrE1 organoid and BHPrS1 spheroid culture was performed using Matrigel (Corning, 356230) based on the manufacturer’s instruction and previously published protocols (50–52) with minor modifications. Briefly, 700 cells/well of BHPrE1, or 14000 cells/well of BHPrE1, or 700 BHPrE1 cells plus 1400 BHPrS1cells per well for co-culture were suspended in 10 µL of DMEM/F-12 medium and then mixed with 290 µL of 5 µg/ml of Matrigel. The 300 µL of cell-Matrigel mixture was then added to a 24-well plate with 100 µL pre-casted Matrigel marix base in each well. After matrix gel formed, organoid cultures were maintained in 500 μL of DMEM/F-12 media that was changed every 3-4 days.

The organoids and spheroids were imaged under Nikon Eclipse TS2 microscope with CaptaVision+ software. The BHPrS1 spheroid colonies were treated with 100 μM Finasteride or DMSO for 72 hours after 14 days and collected after 17 days following a previously published protocol (52, 53). The fixed spheroid colonies were analyzed with immunofluorescence for Ki-67 and LEF1.

### Softy Agar Colony Formation Assay

BPH-1 growth in different conditioned mediums was determined by assaying colony formation in soft agar (54). Briefly, 5,000 BPH-1 cells suspended in 2x RPMI1640 (Gibco, 31800022) with 20% FBS were mixed with 0.6% Noble Agarose (RPI, A20460) and then seeded on top of 1% noble agarose layer in triplicates in 6-well plates. The cells were topped with 500uL condition media and cultured for 21 days in a humidified incubator containing 5% CO_2_ at 37 °C with media replacement every 3-4 days. Colonies formed were then stained with 1mg/mL crystal violet in 1xPBS with 10% methanol overnight. Images of colonies were taken under a white view using Nikon Eclipse TS2 microscope and CaptaVision+ software.

### Cell viability assay

The proliferation of BHPrS1 stromal cells were measured with MTT assay (Roche, 11465007001) as previously described (55), Briefly, 1000 cells/well of BHPrS1 cells in 200 µL/well of culture medium were seeded in 96-well plates and then incubated for 72 hours with corresponding treatments. The proliferation of cells was determined by the relative absorbance reading of 590nM measured using SpectraMax id5 plate reader.

### Immunocytochemistry (ICC)

Cells seeded on coated coverslips (Electron Microscopy Sciences, 7229011) in 12-well plates were fixed in 4% paraformaldehyde (Electron Microscopy Sciences, 15710) and non-specific binding of antibodies was blocked using 3% BSA (Boston Bioproducts, P-753) in 1x PBS. Fixed cells were permeabilized with 0.1% Triton X-100 (MP Biomedical, 04807426) in 1xPBS (PBSX) and incubated with primary antibody diluted in 3% BSA in PBSX for 90 minutes followed up with corresponding secondary antibodies for 60 minutes at room temperature. After excess antibodies washed away, samples were mounted with antifade reagent with DAPI (Invitrogen, S36939). For quantitative analysis, the fluorescence of the corresponding antibodies was imaged with Keyence BZ-X 2000 microscope and the intensity was measured with Image J. All antibodies used in this study were obtained from various sources and listed in Supp. Table S1.

### Immunohistochemistry (IHC)

IHC staining on MTOPS biopsy samples and BIDMC biorepository surgical samples were performed as previously described (14). Briefly, fixed prostate tissue sections undergo deparaffinization, antigen retrieval, blocking, antibody application (detailed in Supp. Table S1), and counterstaining for visualizing SRD5A2 and WNT5A. Probed sections were imaged and average immunoreactive score (IRS) was calculated following the same protocol established in the previous publication (14).

### Immunofluorescence (IF)

IF staining on BHPrS1 spheroid-like colonies was performed similarly to ICC described in previous subsection with minor modifications. Briefly, cell colonies were collected and fixed on coverslips in 12-well plates with 4% PFA. Rinsed plates were frozen at −20 degrees for storage. Upon thawing, the colonies on coverslips were blocked, permeabilized, probed with corresponding antibodies (detailed in Supp. Table S1), and imaged following the ICC protocol.

### Cell Lysate Preparation and Western Blot

Cells cultured in 6-well plates were washed with sterile 1XPBS (Cyvita, SH30256.FS) and detached with trypsin (Gibco, 2520114). Detached cells were then pelleted via centrifugation and subsequently frozen at −80°C. Upon thawing, cells were lysed in 20 volume size of 1x M-PER buffer (ThermoFisher, 78501) with 0.1% SDS (Boston BioProducts, BM-230) and 1% protein and protease inhibitor cocktail (ThermoFisher, 78445), and chilled on ice for 30 minutes with constant shaking. Insoluble cell debris was removed after centrifugation at 13,200g for 15 minutes at 4°C, and the protein concentration of the clear supernatant was measured with the BCA assay (ThermoFisher, 23227).

Western blot was performed as previously described (36) with minor modifications. Cell lysate samples were diluted with lysis buffer to reach the same protein concentration and then denatured in 6x Laemmli Buffer (AlfaAesar, J61337) with 10% 2-Mercaptoethanol (SigmaAldrich, M3148) for 5 minutes at 95°C. Denatured protein samples were run on 8-16% SDS-PAGE gels (Bio-rad, 4568104) at 90 volts for 90 minutes and subsequently transferred to PVDF membrane (Millipore, IPVH00010) for 3 hours on 250 mA at 4°C. The membrane was blocked with 5% (w/v) non-fat dry milk (Labscientific, M0841) in 1x TBS buffer (diluted from 20x TBS buffer, Boston Bioproducts, BM-301X), followed by probing with corresponding antibodies diluted in 5% BSA in 1x TBST buffer (1xTBS buffer with 0.05% Tween-20, FisherScientific, BP337-500). Probed membrane were developed using ECL reagents (Cyvita, RPN2232) and imaged with LI-COR Odyssey M imager. Antibodies used in this study were obtained from various sources and listed in Supp. Table S1.

### Quantitative PCR (qPCR) and PCR Microarray

qPCRs were performed as previously described (36) and run on Applied Biosystems QuantStudio 6 Flex. GraphPad Prism 10 was utilized to calculate the ratio of mRNA levels of assayed genes to values of GAPDH control using the ΔC_T_ method (2^−ΔΔCT^). Primers were ordered from IDT and listed in Supp. Table S2.

The WNT signaling pathway RT^2^ PCR Microarray was performed using Human WNT signaling pathway RT^2^ profiler PCR array (Qiagen, 330231 PAHS-243ZA) and corresponding reverse transcription kit (Qiagen, 330401) and RT2 SYBR green qPCR master mix (Qiagen, 330500) following the manufacturer’s protocol.

### ELISA

WNT5A and IGF1 level in cell and culturing medium were quantified using human WNT5A ELISA kit (VWR, ANTIA77507-96) or IGF1 ELISA kit (SigmaAldrich, RAB0228) following the manufacturer’s protocol. The absolute concentration was calculated by plotting the mean absorbance values of the unknowns onto the standard curve in GraphPad Prism 10, and the cellular concentration was calculated by dividing the value of absolute concentration over the protein concentration of corresponding cell lysate samples.

The DHT level in the BIDMC biorepository serum samples and frozen surgical prostate tissues were measured with human DHT ELISA kit (LS Bio, LS-F10533-1) following the manufacturer’s standard protocol as previously described (14).

### Single-cell RNA sequencing (scRNA-seq)

ScRNA-seq and data analysis was performed as previously described (14). Briefly, 23,000 single cells from fresh prostate tissues were loaded onto separate lanes of a single-cell A Chip and generated the libraries submitted for sequencing on an Illumina NextSeq 500 and NovaSeq 600 sequencers.

### RNA Fluorescence *in situ* Hybridization (RNA-FISH)

RNA-FISH analysis of prostate tissue was performed as previously described (14). Briefly, three of each age-matched *Srd5a2*-null, heterozygous, and wild-type mice were sacrificed and whole prostate tissues were fixed and sliced for staining conducted at the Neurobiology Imaging Facility, Harvard Medical School. Image acquisition and quantitative analysis of *Wnt5a* signals follows the same protocol established in previous publication (14).

### Statistical Analysis

All data included in this study were presented as mean ± standard deviation with at least three independent experiments. A *P* value of equal or less than 0.05 was considered significant. The statistical methods for each corresponding experiment are described in the figure legends.

### Data and Resource Availability

Further information, results, reagents, and other supporting data in this study are available from the corresponding author upon request. Raw data is available in the supplemental files.

## Acknowledgements

We thank Dr. Chad Vezina’s laboratory at the University of Wisconsin-Madison for generously providing us with Srd5a2^+/–^ animals for breeding. Dr. Linus Tsai at BIDMC conducted scRNA-seq at the Boston Nutrition and Obesity Center Functional Genomics and Bioinformatics Core. The MTOPS study was conducted by the MTOPS study investigators and supported by the NIDDK. The data and biospecimen from the MTOPS study reported here were supplied by the NIDDK Central Repository. This manuscript was not prepared in collaboration with investigators of the MTOPS study and does not necessarily reflect the opinions or views of the MTOPS study, the NIDDK Central Repository, or the NIDDK.

## Authors’ contributions

ZW and AO performed study design, conceptualization, and supervised this research. ZW and CS designed, executed, and interpreted the experiments on mouse models. BG designed, executed, and interpreted molecular biology, cell biology, and *in vitro* experiments. XL provided bioinformatics analysis. CS, NP, SL and YT conducted and interpreted IHC experiments and provided reagents. CS, BG, and ZW analyzed data, designed figures, and wrote the manuscript. All authors reviewed and approved the manuscript.

## Ethics Approval and Consent to Participate

Studies on human tissues from MTOPS and institutional cohort were performed on de-identified patient samples, and these studies were deemed exempt by the Institutional Review Board at Beth Israel Deaconess Medical Center.

All animal work described in this study was **i**n adherence to US NIH Guide for the Care and Use of Laboratory Animals and approved by the Beth Israel Deaconess Medical Center Institutional Animal Care and Use Committee (protocol number 049-2022).

## Data Availability Statement

The scRNA-seq data are available in NCBI’s GEO (https://www.ncbi.nlm.nih.gov/geo) under accession numbers GSM7504455 (Mm-prostate-02), GSM7504456 (Mm-prostate-04), GSM7504457 (Mm-prostate-06) under previous publication (14). Raw data of quantifications in figures is available in the supplemental Supporting Data Values file. Uncropped and unprocessed Western blots are provided in Supplemental Unedited Blot Images file. Further information, results, reagents, and other supporting data used or analyzed in this study are available from the corresponding author upon request.

The authors declared no conflict of interest.

