## Supplementary figures and images for "Stromal SRD5A2 promotes prostate growth through WNT5A-LEF1-IGF1 signaling in benign prostatic hyperplasia"

### Supplemental Figure 1

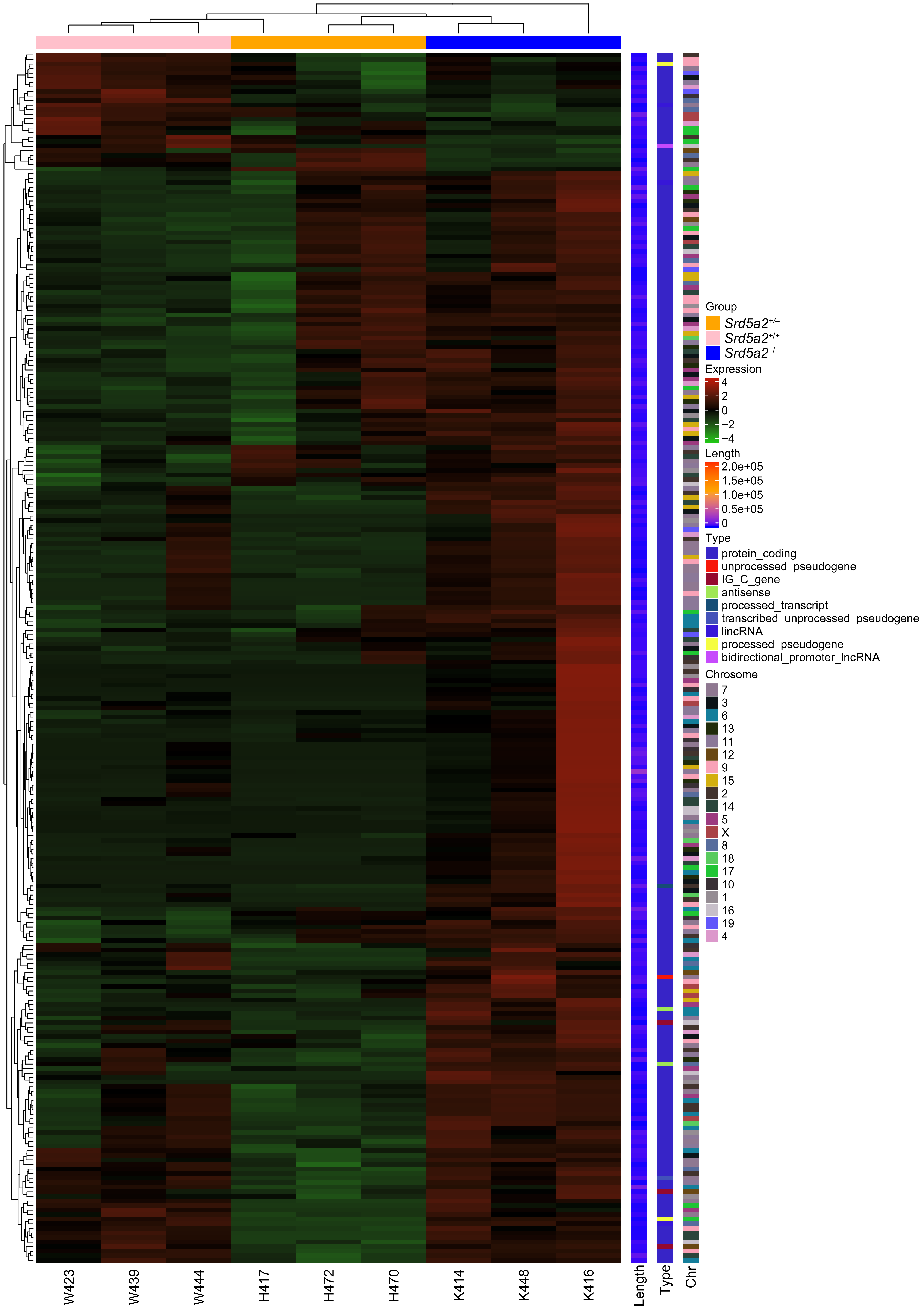

### Supplemental Figure 2

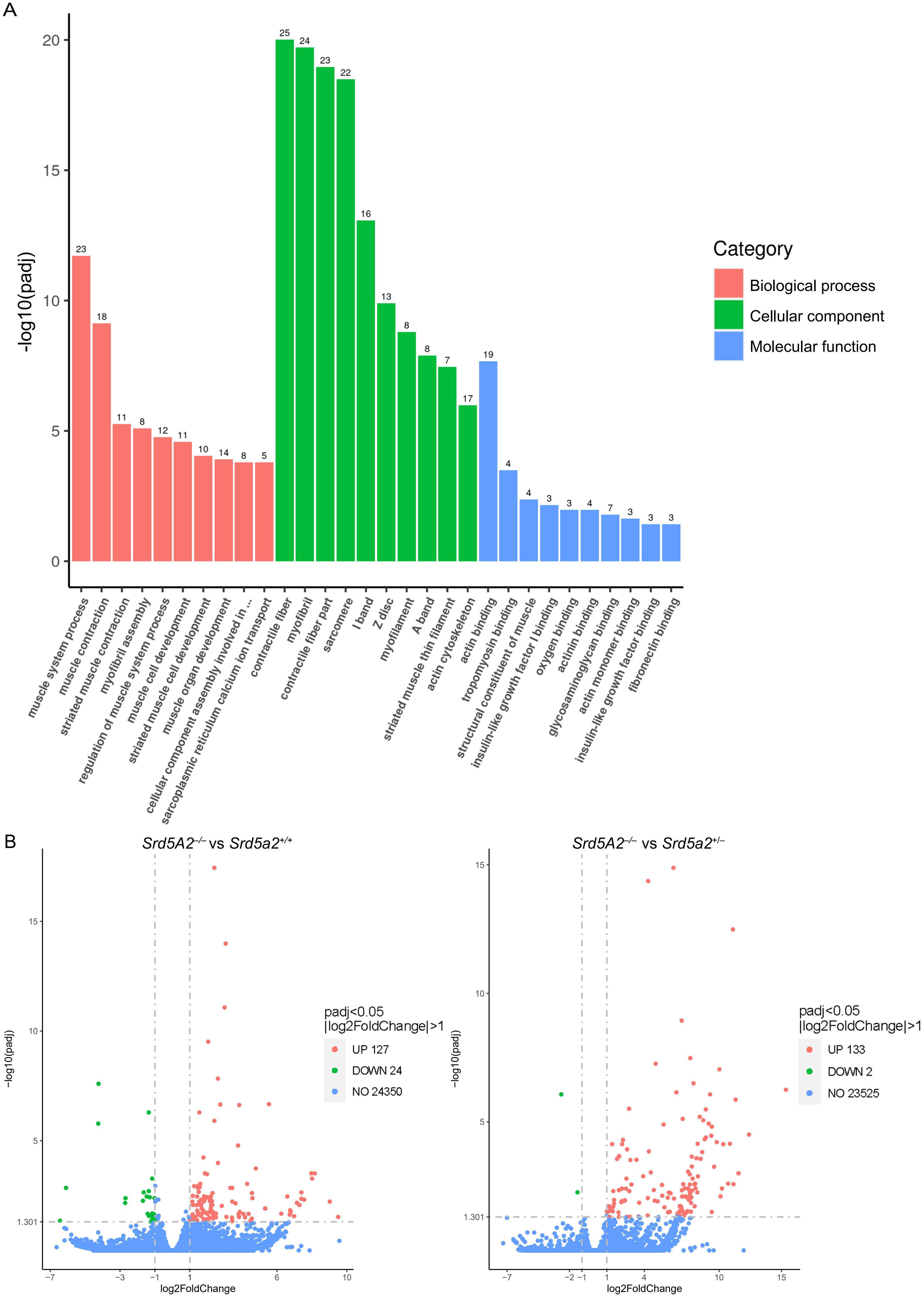

### Supplemental Figure 3

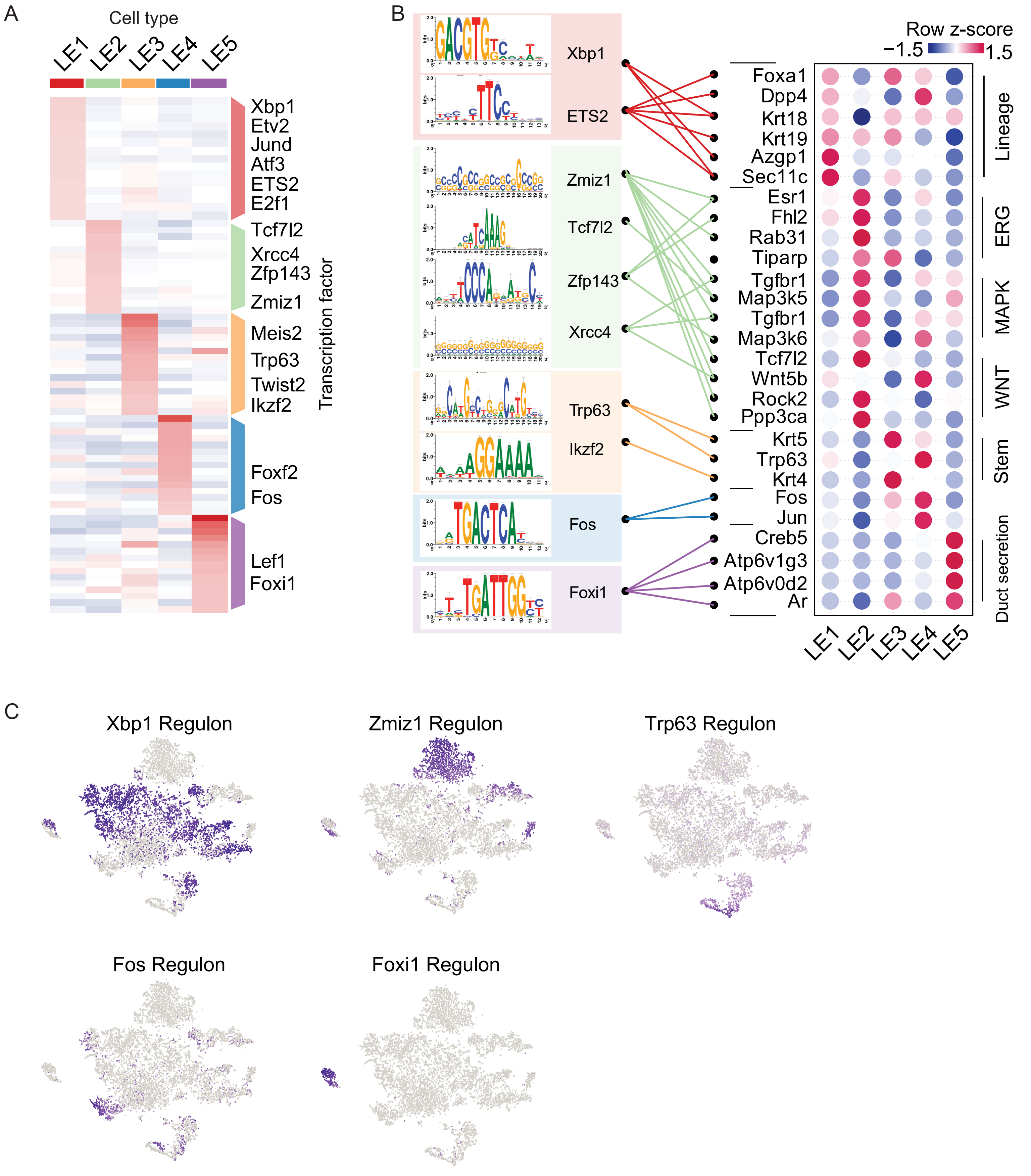

### Supplemental Figure 4

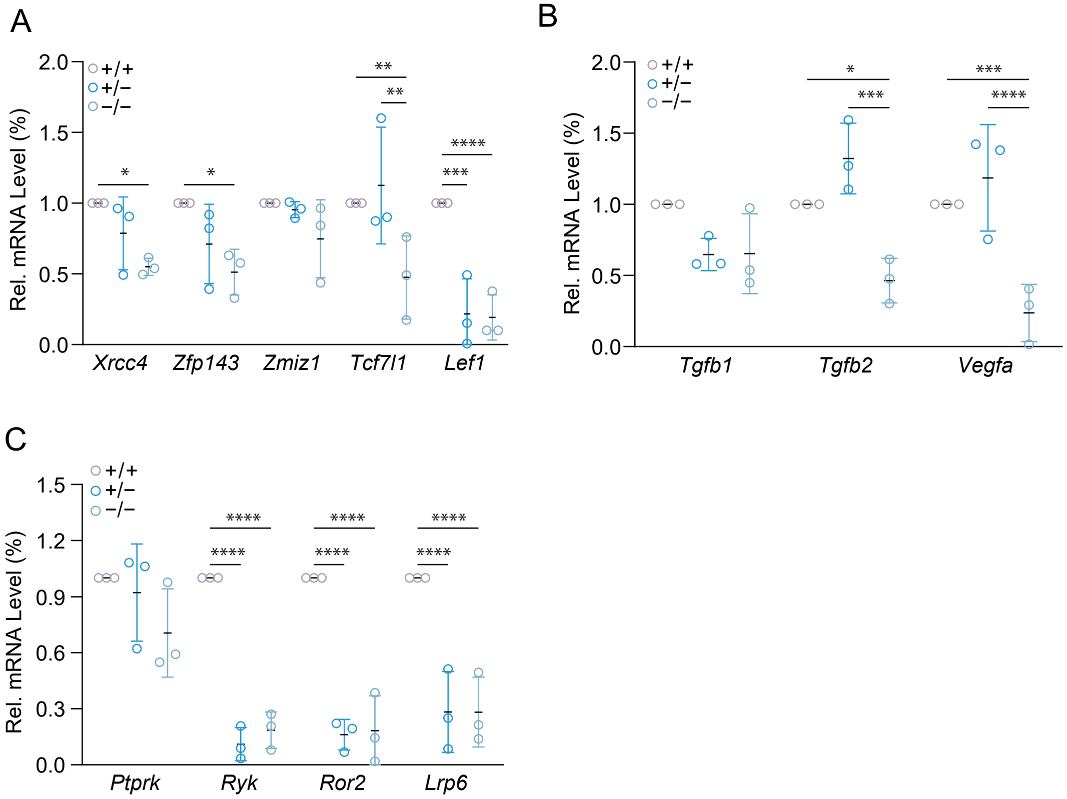

### Supplemental Figure 5

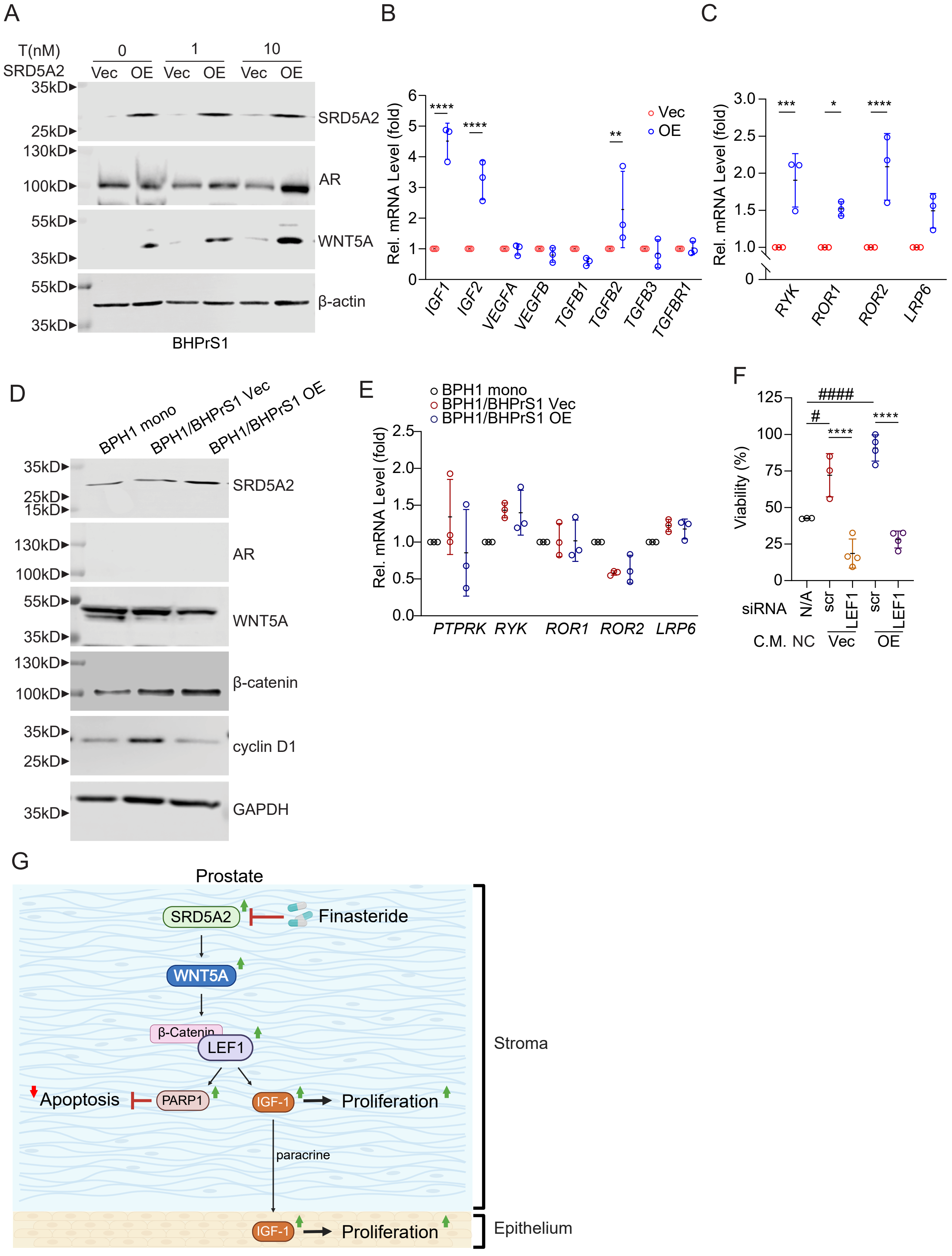
