## Supplemental Figure legends for "Stromal SRD5A2 promotes prostate growth through WNT5A-LEF1-IGF1 signaling in benign prostatic hyperplasia"

**Supplementary Figure 1. Hierarchical clustering and differential gene expression analysis in** ***Srd5a2* genotypes.** To evaluate the molecular changes caused by SRD5A2 KO, bulk RNA-seq was conducted on *Srd5a2^+/+^*, *Srd5a2*^+/-^*, Srd5a2*^-/-^ prostate tissues to identify putative pathways involved in these processes. Hierarchical clustering successfully separated these groups based on differential gene expression. Significantly upregulated (133) and downregulated (2) genes for KO mice were determined using gene ontology (GO) analysis with Fisher’s exact test.

**Supplementary Figure 2. Gene ontology (GO) enrichment analysis of Differentially Expressed Genes (DEG) in *Srd5a2* genotypes.** Bulk RNA sequencing was performed on prostate tissues from *Srd5a2*^+/+^, *Srd5a2*^+/−^, and *Srd5a2*^−/−^ mice. **(A)** GO enrichment analysis was performed on significantly differentially expressed genes between *Srd5a2^+/+^* and *Srd5a2*^-/-^prostate tissues. The top enriched GO terms are categorized into biological processes (BP, red), cellular components (CC, green), and molecular functions (MF, blue). The x-axis shows the GO terms, while the y-axis represents the -log10 (adjusted p-value) of enrichment significance. The numbers above the bars indicate the count of genes associated with each GO term. **(B)** DEG analysis between *Srd5a2*^+/+^, *Srd5a2*^+/−^, and *Srd5a2*^−/−^ mice prostate tissues. Left: A volcano plot illustrates significantly upregulated (n=127) and downregulated (n=24) genes in the *Srd5a2*^-/-^  mice compared to *Srd5a2^+/+^* controls, using a threshold of log₂FoldChange >1 and adjusted p-value <0.05. Right: A volcano plot illustrates significantly upregulated (n=133) and downregulated (n=2) genes in the *Srd5a2*^-/-^  mice compared to *Srd5a2^+/-^* controls.

**Supplementary Figure 3. Transcription factor regulatory networks and gene expression signatures across luminal epithelial subtypes in mouse prostate. (A)** Heatmap displaying the activity of transcription factors across five luminal epithelial (LE1–LE5) clusters identified from scRNA-seq. Row z-scores indicate the relative activity of key transcription factors in each cluster. **(B)** Motif logos and regulatory networks of selected transcription factors linked to their target genes. Colored edges represent different regulon groups associated with luminal epithelial identity. Dot plot showing expression patterns of representative lineage, ERG signaling, MAPK signaling, WNT signaling, stemness, and duct secretion genes across LE1–LE5 clusters. The color represents expression level (row z-score), and the size of the dot reflects the proportion of expressing cells within each cluster. **(C)** Spatial distribution of key regulon activities across mouse prostate cells. UMAP plots showing the spatial distribution of regulon activity for selected transcription factors. Cells with higher predicted regulon activity are shown in purple, while cells with low or absent activity are shown in gray. Each plot highlights the distinct cellular localization patterns of transcription factor regulatory networks within the prostate epithelial cell population.

**Supplemental Figure 4. SRD5A2 knockout impaired prostate mouse prostate gene profiles.** (**A-C**) Quantitative PCR analysis of transcription factors (A), secreted ligands (B), and WNT5A receptors (C) identified from scRNA-seq analysis in wildtype (+/+), *Srd5a2* hetero (+/-), and *Srd5a2* null (-/-) mice whole prostate. N≥3 in each condition. Error bar stands for mean ± SD. Student’s t-test and one-way ANOVA were performed. ns, p > 0.05; *, p ≤ 0.05; **, p ≤ 0.01; ***, p≤ 0.001; ****; p ≤ 0.0001.

**Supplemental Figure 5. SRD5A2-WNT5A-LEF1 regulatory axis in prostate cells.** (**A**) Western blot analysis of SRD5A2, androgen receptor (AR), WNT5A, and β-actin in control and SRD5A2-overexpressing BHPrS1 cells. (**B-C**) Quantitative PCR analysis of WNT-related growth factors (B) and WNT5A receptors (C) in control and SRD5A2-overexpressing BHPrS1 cells. (**D**) Western blot analysis of SRD5A2, AR, WNT5A, β-catenin, cyclin D1, and GAPDH loading control in BPH1 cells co-cultured with BPH1 cells, BHPrS1 control cells, or BHPrS1 SRD5A2 overexpressing cells. (**E**) Quantitative PCR analysis of WNT5A receptors in BPH1 cells co-cultured with BPH1 cells, BHPrS1 control cells, or BHPrS1 SRD5A2 overexpressing cells. (**F**) Cell viability of BPH1 cells cultured in siRNA-transfected BHPrS1-conditioned medium for 72 hours measured by MTT assay. (**G**) Proposed schematic model of SRD5A2-regulated WNT5A expression and SRD5A2 regulating prostate growth via PARP1 and IGF-1. N≥3 in each condition. Error bar stands for mean ± SD. Student’s t-test and one-way ANOVA with Tukey test were performed. *, p ≤ 0.05; **, p ≤ 0.01; ***, p≤ 0.001; ****; p ≤ 0.0001. #, significance between different groups with the same meaning as *.
